# Mitochondrial mRNA localization is governed by translation kinetics and spatial transport

**DOI:** 10.1101/2022.02.16.480701

**Authors:** Ximena G Arceo, Elena F Koslover, Brian M Zid, Aidan I Brown

## Abstract

For many nuclear-encoded mitochondrial genes, mRNA localizes to the mitochondrial surface co-translationally, aided by the association of a mitochondrial targeting sequence (MTS) on the nascent peptide with the mitochondrial import complex. For a subset of these co-translationally localized mRNAs, their localization is dependent on the metabolic state of the cell, while others are constitutively localized. To explore the differences between these two mRNA types we developed a stochastic, quantitative model for MTS-mediated mRNA localization to mitochondria in yeast cells. This model includes translation, applying gene-specific kinetics derived from experimental data; and diffusion in the cytosol. Even though both mRNA types are co-translationally localized we found that the steady state number, or density, of ribosomes along an mRNA was insufficient to differentiate the two mRNA types. Instead, conditionally-localized mRNAs have faster translation kinetics which modulate localization in combination with changes to diffusive search kinetics across metabolic states. Our model also suggests that the MTS requires a maturation time to become competent to bind mitochondria. Our work indicates that yeast cells can regulate mRNA localization to mitochondria by controlling mitochondrial volume fraction (influencing diffusive search times) and gene translation kinetics (adjusting mRNA binding competence) without the need for mRNA-specifc binding proteins. These results shed light on both global and gene-specific mechanisms that enable cells to alter mRNA localization in response to changing metabolic conditions.

## Introduction

To sustain life and function, cells maintain a homeostatic internal state while retaining the capacity to respond to variable environments and challenges. For eukaryotic cells, homeostasis requires not only regulation of gene expression, but also maintainance of internal organization through the sorting of proteins among organelles and subcellular compartments. Spatial targeting of proteins to specific cellular destinations can occur through a variety of transport and retention mechanisms, sometimes acting in combination (***Bauer et al., 2015***; ***Aviram and Schuldiner, 2017***; ***Chio et al., 2017***; ***Guardia et al., 2018***; ***Wheeler and Hyman, 2018***; ***Maza et al., 2019***; ***Mogre et al., 2020***).

Protein localization is often controlled by first transporting the mRNA to a specific region (***Das et al., 2021***), and then translating proteins locally. mRNA localization serves as a key mechanism for delivering proteins to far-flung cell regions in neurons (***Biever et al., 2019***), expediting protein synthesis when locally required (***Bramham, 2008***), and ensuring proteins are provided a suitable environment for folding (***Stephens et al., 2005***). Failure to localize mRNA can result in developmental defects (***Trcek and Lehmann, 2019***) and cognitive disorders (***Das et al., 2019***).

Canonical descriptions of protein localization through mRNA transport include translational suppression en route (***Besse and Ephrussi, 2008***; ***Das et al., 2021***), with protein synthesis beginning only after the mRNA reaches its target destination. By contrast, some mRNA are known to begin translation while in transit (***Wang et al., 2016***; ***Cioni et al., 2019***). For such cases, we explore how translational dynamics themselves can control mRNA localization, focusing on nuclear-encoded mitochondrial genes in yeast.

While some mitochondrial genes are encoded by mitochondrial DNA, the vast majority of mitochondrial proteins are translated from nuclear-encoded mRNA (***Boengler et al., 2011***) and a subset of those mRNAs have been observed to localize to the mitochondrial surface. In *Saccharomyces cerevisiae* these mitochondrially localized mRNAs have been subclassified based on their mechanism of localization. Class I mRNAs are primarily targeted to the mitochondria by the RNA binding protein Puf3, while Class II mRNAs localize independently of Puf3 (***Devaux et al., 2010***; ***Saint-Georges et al., 2008***). Class II mRNAs are proposed to localize through translation of the amino-terminal mitochondrial targeting sequence (MTS) that can associate with import complexes on the cytosolic side of the outer mitochondrial membrane (***Garcia et al., 2010***).

*S. cerevisiae* yeast rely heavily on glucose fermentation even in aerobic conditions. With non-fermentable carbon sources, the shift to a respiratory metabolism involves dramatic changes to the mitochondrial proteome (***Di Bartolomeo et al., 2020***; ***Morgenstern et al., 2017***). This shift also leads to an increase in the fraction of the cytosol occupied by mitochondria (mitochondrial volume fraction, or MVF) (***Tsuboi et al., 2020b***), which form dynamic tubular networks distributed throughout the cell (***Viana et al., 2020***). While Class II mRNAs were initially found to be mitochondrially localized under respiratory conditions, many exhibit condition-dependent localization, as almost 70% do not robustly localize to mitochondria under fermentative conditions (***Williams et al., 2014***; ***Tsuboi et al., 2020b***,a). This may be due at least in part to changes in MVF, which can quantitatively predict the conditional localization behavior of mRNAs *ATP2* and *ATP3* (***Tsuboi et al., 2020b***). Additionally, many Class II mRNAs that do not robustly localize under fermentative conditions, including *ATP2* and *ATP3*, become mitochondrially localized upon application of the translation elongation inhibitor cycloheximide (CHX) (***Williams et al., 2014***; ***Tsuboi et al., 2020b***). By contrast, other Class II mRNAs such as *TIM50* have high, constitutive localization to mitochondria even in fermentative conditions (***Tsuboi et al., 2020b***), and respond little to increased MVF (***Tsuboi et al., 2020b***) or CHX application (***Williams et al., 2014***). Given that all Class II mRNAs contain an MTS but only some are localized under fermentative conditions, these observations suggest that the presence of the MTS is required but not sufficient for preferential localization to mitochondria. This idea has been further supported through MTS swapping experiments (***Garcia et al., 2010***).

Localization of a Class II mRNA to a mitochondrion requires exposure of an MTS peptide sequence while the mRNA is very near to the mitochondrial membrane, implying that such localization can be modulated through the relative kinetics of MTS exposure and spatial movement throughout the cell. By arresting translation, CHX leaves nascent peptides and any of their translated MTS motifs exposed indefinitely. The increase in mRNA localization upon CHX application thus substantiates the importance of gene-specific translation dynamics for mitochondrial localization. Similarly, the dependence of mitochondrial localization on the MVF suggests that the geometry encountered by a diffusing mRNA can meaningfully control the frequency of mitochondrial proximity and opportunities for an MTS to interact with a mitochondrial surface.

The physical process of localization requires a transport mechanism enabling an mRNA to encounter its target region and a retention mechanism to limit mRNA escape. In the relatively small volume of a yeast cell, diffusion is sufficient to distribute mRNA, with diffusive arrival rates to cellular targets modulated by intracellular geometry (***Saffman and Delbrück, 1975***; ***Berg and Purcell, 1977***; ***Reguera and Rubi, 2001***; ***Condamin et al., 2007***; ***Koslover et al., 2011***; ***Brown et al., 2020***; ***Mogre et al., 2020***). Once an mRNA has diffusively reached a destination, binding interactions then determine the time period of mRNA localization. Equilibrium mRNA localization would be determined by the probability of occupying a binding-competent state and the volume of the localization region, i.e. the MVF. However, the energy-consuming process of translation pushes mRNA localization out of equilibrium, similar to other driven processes necessary to maintain cellular organization, including protein targeting (***Murugan et al., 2012***; ***Gladrow et al., 2016***; ***Maza et al., 2019***; ***Brown and Sivak, 2019***; ***Fang and Wang, 2020***; ***Mogre et al., 2020***).

To address how translational dynamics could control the localization of mRNA for mitochondrial genes, we developed a stochastic, quantitative model for mitochondrial mRNA localization that incorporates translation and diffusion within a yeast cell. The model is parameterized against published genome-wide measurements of both constitutively and conditionally localized Class II mRNAs (***Couvillion et al., 2016***; ***Morgenstern et al., 2017***; ***Zid and O’Shea, 2014***). We find that the kinetics of translation, as well as the diffusive search time-scales, determine the level of mRNA localization to mitochondria, enabling both low and high localization within the physiological range of key parameters. Crucial to our description of mitochondrial mRNA localization is a proposal for an MTS maturation time following translation of the MTS peptide sequence. Our work suggests a distinct mode of spatial protein regulation and a mechanism for yeast and other cells to control protein localization using gene-specific translation dynamics combined with global adjustments of organelle size.

## Results

### Localization depends on both equilibrium and kinetic contributions

To help guide our investigation of the translational control of mRNA localization, we begin by analyzing a general minimal model (Fig. 1A). We assume that mRNA is capable of switching between a binding-competent (“sticky”) state and a binding-incompetent (“non-sticky”) state. For mitochondrial targeting, a binding-competent state corresponds to an mRNA with at least one partially-translated peptide with an exposed MTS sequence. We define two rate constants: *k*_S_ and *k*_U_ for switching into and out of the competent state, respectively, assumed to be independent of the mRNA location. At equilibrium the fraction *f*_s_ = *k*_S_/(*k*_S_ + *k*_U_) is in the competent state. Fora binding-competent mRNA to bind to a mitochondrion, it must be sufficiently proximal to a mitochondrial surface. Binding-incompetent molecules can move from the bulk into binding range of a mitochondrion with rate *k*_R_ and can leave the near-surface region with rate *k*_L_. These rates are expected to depend on the diffusivity of the mRNA and the geometry (size and shape) of mitochondria within the cell. At equilibrium, *f*_d_ = *k*_R_/(*k*_R_ + *k*_L_) is the fraction of the mRNA-accessible cell volume that is within binding range of the mitochondrial surface. As the cytosolic volume fraction that is near mitochondria, *f*_d_ is distinct from but related to the MVF, the cell volume fraction occupied by mitochondria. The binding-competent mRNA reach the mitochondrial region with the same rate *k*_R_ but are assumed to bind irreversibly and cannot leave until they switch into the incompetent state.

**Figure 1.**
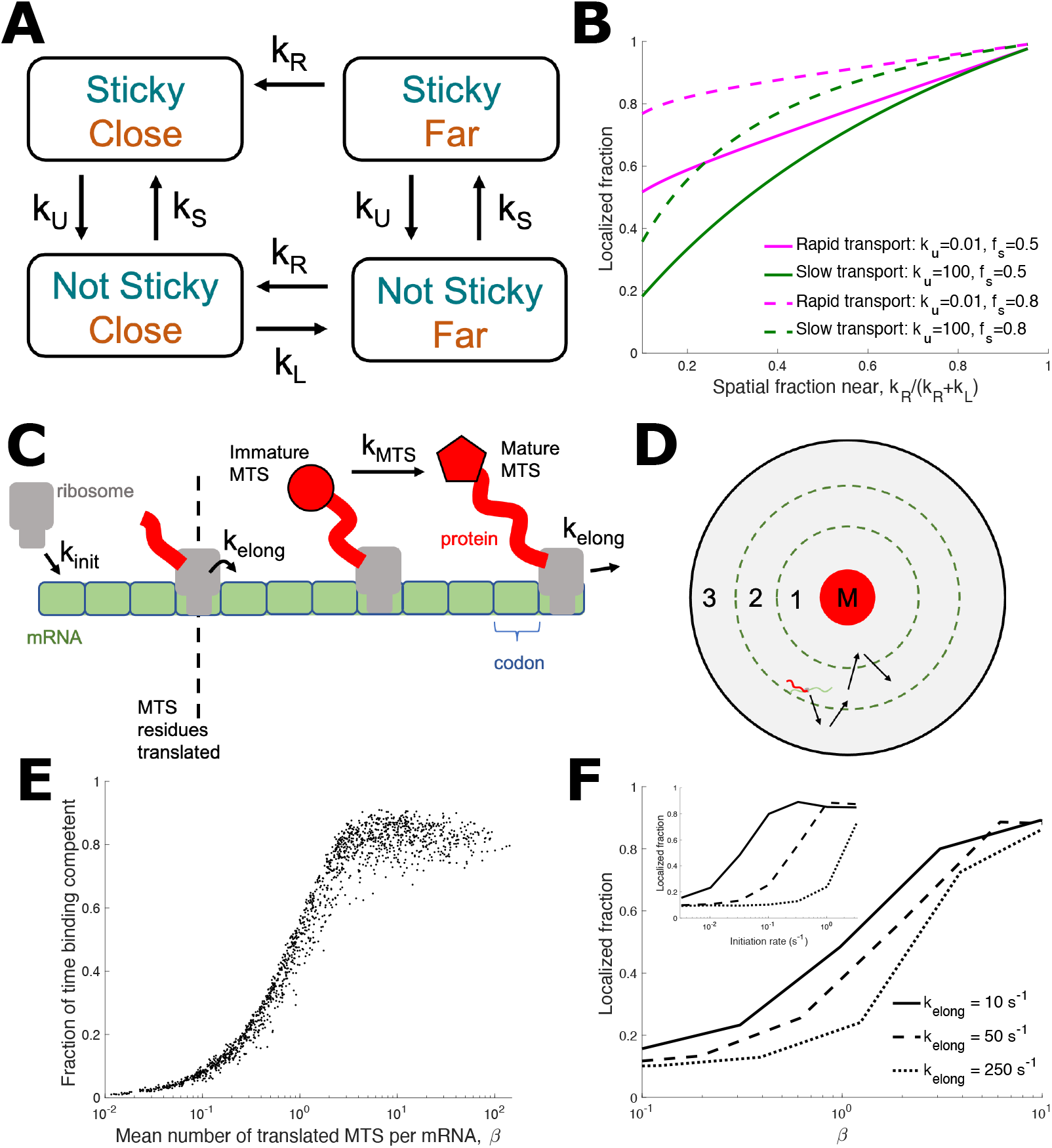
Quantitative models show equilibrium and kinetic contributions to mitochondrial mRNA localization. (A) Simplified discrete-state model of mRNA mitochondrial localization. mRNA can be either binding competent (‘sticky’) or not binding competent (‘not sticky’), and either within binding range of mitochondria (‘close’) or not within binding range (‘far’). mRNA transition between these states with rates described in the text. (B) Localized fraction [defined as ‘close’ in (A)] as the spatial fraction of the cell near mitochondria is varied. Rapid transport curves indicate rapid switching from close to far relative to switching between sticky and not sticky, while for slow transport the relative switching speeds are reversed. (C) Stochastic model of mRNA translation. Ribosomes initiate translation at rate *k*_init_ and progress to the next codon at rate *k*_elong_. MTS is translated after the first 100 amino acids. Once MTS is translated, MTS becomes binding-competent at rate *k*_MTS_. (D) Schematic of mRNA diffusion in spatial model. The cytoplasmic space is treated as a cylinder centered on a mitochondrial cylinder. mRNA in region 1 are sufficiently close for binding-competent mRNA to bind to the mitochondria, mRNA in region 2 are considered mitochondrially localized in diffraction-limited imaging data, and region 3 represents the remainder of the cell volume. mRNA not bound to mitochondria will freely diffuse between these regions. (E) For the stochastic translation model shown in (C), the fraction of mRNA lifetime that an mRNA is binding-competent vs. *β* = *k*_init_(*L* – *L*_MTS_)/*k*_elong_, the mean number of translated MTSs per mRNA. For each data point, mRNA translation parameters *k*_init_, *L*, and *k*_elong_ were randomly selected from the ranges *k*_init_ ∈ [10^-3^ s^-1^, 0.5 s^-1^], *L* ∈ [150 aa, 600 aa], and *k*_elong_ ∈ [1 s^-1^, 10 s^-1^]. (F) Mitochondrial localization from the stochastic model illustrated in C and D, as *β* is indirectly varied through *k*_init_ adjustment. *L* = 400 aa, 4% mitochondrial volume fraction, and *k*_elong_ as indicated in legend. Inset displays the same data, but plotted directly against *k*_init_.

The resulting four-state model (binding-competent vs not, proximal to mitochondria vs not) is illustrated in Fig 1A. Given the assumed irreversible binding of competent mRNAs, the model is inherently out of thermal equilibrium. The kinetic equations can be solved to find the steady-state fraction of mRNA localized to the proximal region, as a function of the kinetic rates (see Methods).

The solutions exhibit two limiting regimes of interest. In the rapid-transport regime where mRNA transport is much faster than the competence switching rate (*k*_U_, *k*_S_ ≪ *k*_R_, *k*_L_), incompetent mRNA can equilibrate throughout the entire cell prior to a switching event. Similarly, competent mRNA can rapidly reach the proximal region and bind to mitochondria. The fraction of mRNA that are mitochondrially localized is then given by the two equilibrium fractions,

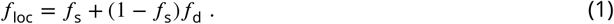

In this spatially equilibrated situation, changing the mitochondrial volume fraction would affect only *f*_d_. If binding dynamics are held fixed (fixed *f*_s_), the mitochondrially localized fraction *f*_loc_ will depend linearly on the proximal volume fraction *f*_d_, with the slope determined by the equilibrium binding competence *f*_s_.

In the opposite slow-transport regime, mRNA transport is much slower than the switching rate (*k*_R_, *k*_L_ ≪ *k*_U_, *k*_S_) and the fraction localized is given by:

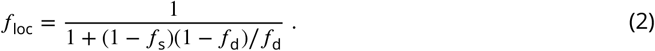

This regime exhibits nonequilibrium behavior. In the limit of low mitochondrial volume fraction (*f*_d_ ≪ 1), the localization probability goes to zero. This is a fundamental difference from the rapid-transport regime, where even at low volume fractions, binding-competent mRNA localize to mitochondria. As a result, the regime with slow transport and fast switching is expected to exhibit a steeper, more non-linear increase in localization with increasing mitochondrial volume fraction (green lines in Fig. 1B) compared to the rapid-transport regime (magenta lines in Fig. 1B.).

This highly simplified, analytically tractable, four-state model is agnostic to the mechanistic details for how the switching between binding-competent and incompetent states occurs, as well as the geometric details of diffusive transport to and from the mitochondria-proximal region. Specifically, it highlights some important non-intuitive features of localization for any molecule that can switch between competent and incompetent states. Namely, the localization behavior is expected to depend not just on the equilibrated binding-competent fraction *f*_s_ and proximal fraction *f*_d_ but also on the relative kinetics of spatial transport and competence switching. In the nonequilibrium regime of fast switching and slow transport, localization becomes non-linearly sensitive to the volume fraction of the target region.

Forthe mitochondrial localization of mRNA, the switching times between competentand incom-petent states are determined by translation kinetics that control exposure duration for attached MTS peptide sequences. The transport kinetics are determined by diffusion timescales towards and away from the mitochondrial surface. We next proceed to develop a more mechanistically detailed model for mitochondrial localization that directly incorporates translation and diffusion.

### Stochastic simulation incorporates translation and diffusive kinetics

The translation kinetics model (Fig. 1C) tracks ribosome number and position. Ribosomes initiate translation on an mRNA with rate *k*_init_, and then proceed along the mRNA codons at elongation rate *k*_elong_. The mRNA is *L* codons in length. The number of codons that must be translated to complete the MTS is set to *l*_MTS_ = 100 to account for an MTS length of up to 70 amino acids and a ribosome exit tunnel length of ~ 30 amino acids (***Bacman et al., 2020***; ***Liutkute et al., 2020***). We begin with an ‘instantaneous’ model, where once translation moves past *l*_MTS_, the mRNA-ribosome complex is assumed to be binding competent until translation completes (*k*_MTS_ → ∞ in Fig. 1C). In subsequent sections we will consider alternative binding-competence models with finite *k*_MTS_.

An mRNA can have multiple MTS-containing nascent peptides if a subsequent ribosome initiates and translates another MTS before the prior translation event is complete. The average number of such binding-competent peptides on a given mRNA is given by

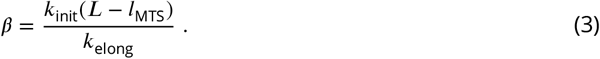

To describe the diffusive encounter of an mRNA with the mitochondrial network, we use a simplified geometric model appropriate for diffusive search towards a narrow tubular target. Specifically, we represent the geometry as a sequence of concentric cylinders (Fig. 1D). The innermost cylinder represents a mitochondrial tubule and serves as a reflective boundary for the mRNA. A slightly larger cylinder represents the region where a binding-competent mRNA is sufficiently close to bind to the mitochondrial surface. If one or more binding-competent MTSs are exposed on an mRNA when it reaches the vicinity of the innermost cylinder, the mRNA will remain associated to the mitochondrial surface until the mRNA returns to zero binding-competent MTSs after peptide translation is completed. A still wider cylindrical region represents locations where the transcript would appear close to the mitochondrial tube in diffraction-limited imaging data, but may not be sufficiently close to bind the mitochondrial surface. Finally, the outermost reflecting cylinder represents the cytoplasmic space available to the diffusing mRNA. The radius of this external cylinder is set such that the innermost mitochondrial cylinder encloses the correct volume fraction of mitochondria to correspond to experimental measurements (which can range from 1%-15%).

This simplified geometry gives an approximate description of the search process for the mitochondrial surface, based on the idea that whenever the mRNA wanders far from any given mi-tochondrial tubule it will approach another tubule in the network, so that its movement can be treated as confinement within an effective reflecting cylinder. Such an approach has previously been used successfully to approximate the diffusive process of proteins searching for binding sites on long coils *of DNA* (***Koslover et al., 2011***). More detailed geometrical features, such as the specific junction distribution and confinement of the yeast mitochondrial network to the cell surface are neglected in favor of a maximally simple model that nevertheless incorporates the key parameters of mitochondrial volume fraction and approximate diffusive encounter time-scale.

Simulations of our stochastic model for simultaneous translation and diffusion can be carried out with any given set of gene-specific translation parameters (*k*_init_, *k*_elong_, *L*). The simulated mRNA trajectories are then analyzed to identify the fraction of mRNA found within the region proximal to the mitochondrial surface (see Methods for details). We find that, for any set of translation parameters, the number of binding-competent MTS sequences (*β*, see Eq. 3) is predictive of the fraction of time (*f*_s_) that each mRNA spends in the binding competent state (Fig. 1E). The greatest variation is near *β* ≈ 1, where different parameter combinations with the same average number of exposed MTSs can give competency fractions ranging from 30 – 50%.

Our analytically tractable 4-state model (Fig. 1B) indicates that localization fraction should depend not only on the binding competent fraction *f*_s_ (related to *β*) but also on the kinetics of switching between competent and incompetent states. We explore the effect of translation kinetics on localization in the stochastic model by varying the initiation and elongation rates of a fixed-length mRNA (Fig. 1F, inset). This approach samples the scope of localization behaviors by simulating multiple combinations of translation parameters. As expected, faster elongation rates (which decrease the period an MTS is exposed on an mRNA and decrease *β*) result in lower localization, and higher initiation rates (which increase *β* while leaving MTS exposure time unaffected) result in higher localization (Fig. 1F, inset). While the number of exposed MTSs, *β*, can explain much of the effect of changing elongation and initiation rates (Fig. 1F), there is substantial variability in localization around *β* ≈ 1, with faster elongation decreasing localization. This result is consistent with the prediction of the 4-state model that rapid switching of binding competence can lead to lower localization even for equal binding competent fractions *f*_s_.

### Physiological translation parameters lead to high mitochondrial binding competence and localization

Because translation kinetics and length vary between genes, we expect the kinetics of binding-competence switching and thus the mitochondrial localization to be gene-specific. To explore the relationship between translation kinetics and mitochondrial localization, we define two categories of Class II mRNAs that were all found to be localized in respiratory conditions (***Saint-Georges et al., 2008***) by their localization sensitivity to translation elongation inhibition by cycloheximide (CHX) in fermentative conditions (***Williams et al., 2014***). “Constitutive” mRNAs preferentially localize to mitochondria both in the absence and presence of CHX. “Conditional” mRNAs do not preferentially localize to mitochondria in the absence of CHX, but do so following CHX application.

Using protein per mRNA and ribosome occupancy data, we estimated the gene specific initiation rate *k*_init_ and elongation rate *k*_elong_ for 52 conditional and 70 constitutive genes (see Methods). Along with the known mRNA lengths *L*, these parameters quantitatively describe translation of each gene in the yeast transcriptome. These measurements indicate that conditional and constitutive genes have similar distributions of ribosome occupancy (Fig. 2A, inset; see Fig. S1 for similar distributions of conditional and constitutive gene ribosome occupancy derived from (***Arava et al., 2003***)). Conditional and constitutive genes also have similar distributions of the number of exposed MTSs, *β*, as calculated from estimated translation parameters (Fig. 2A). Notably, the predicted *β* values were relatively large, with 90% of both constitutive and conditional mRNA estimated to have *β >* 2. Consequently, the stochastic simulation predicts median localization fractions above 80% for both the conditional and constitutive gene groups, with no significant difference between the two groups (Fig. 2B). Comparison of two specific genes (*ATP3* and *TIM50*) known to have mitochondrial localizations with distinct dependence on mitochondrial volume fraction (***Tsuboi et al., 2020b***) also yielded similarly high localization fractions in stochastic simulations, across all mitochondrial volume fractions (Fig. 2C).

**Figure 2.**
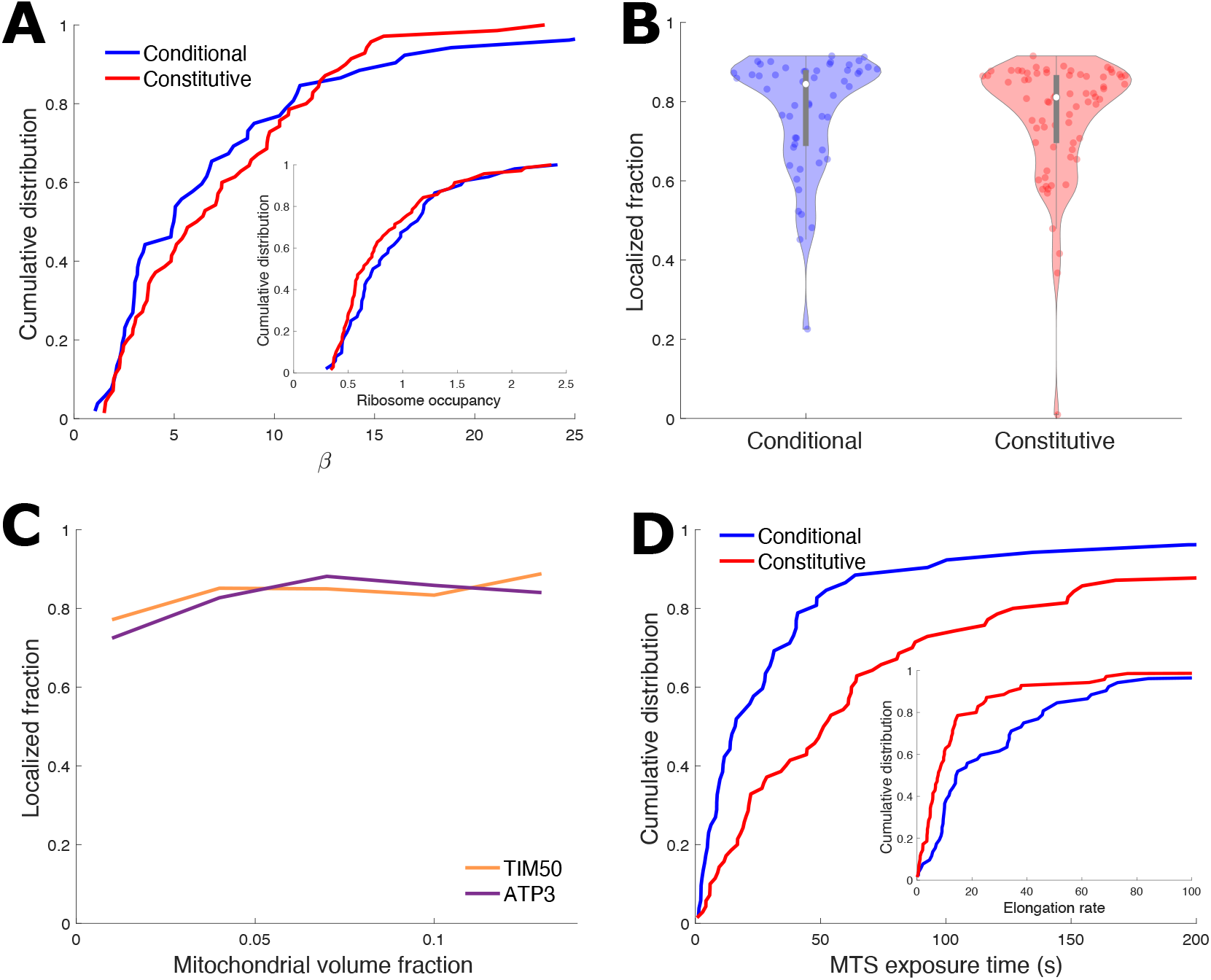
Instantaneous model is insufficient to explain differential mitochondrial localization of different gene groups. (A) Cumulative distributions of conditional and constitutive mRNA genes vs number of binding-competent ribosomes *β* (lines indicate fraction of genes with given *β* or less). *β* for each mRNA gene is calculated from gene-specific *k*_init_ and *k*_elong_ that are estimated from experimental data (see Methods). Inset is cumulative distribution of ribosome occupancy, showing ribosome occupancy and *β* have similar distributions. (B) Violin plot (***Bechtold, 2016***) showing mRNA localization fraction of individual genes with instantaneous model (no maturation delay), with translation kinetics for each gene estimated from experimental data (see Methods). 4% MVF. For direct comparison to experimental data, mRNA in region 1 (see Fig. 1D) recorded as mitochondrially localized. (C) Mitochondrial localization vs mitochondrial volume fraction for *TIM50* and *ATP3* with instantaneous model, with translation kinetics for both genes estimated from experimental data (see Methods). For direct comparison to experimental data, mRNA in regions 1 and 2 (see Fig. 1D) recorded as mitochondrially localized. (D) Cumulative distributions of MTS exposure time *t*_expo_ = (*L* – *l*_MTS_)/*k*_elong_. The steeper rise of conditional genes indicates more conditional gene mRNAs have low exposure times. Translation kinetics for each gene estimated from experimental data (see Methods). Inset shows the cumulative distribution of elongation rate, for which constitutive genes have a steeper rise, indicating slower typical elongation, which contributes to the longer exposure times in the main plot.

These simulation results using gene-specific estimates of the translation parameters *k*_init_, *k*_elong_, and *L* (Fig. 2B,C) run directly counter to experimental measurements. Specifically, they over-predict mitochondrial localization for transcripts, such as *ATP3*, that are known to exhibit low localization values at low mitochondrial volume fractions. Given the high calculated values of *β*, and the importance of MTS exposure kinetics in predicting localization at intermediate *β* values, we more closely examined the quantities underlying this parameter, which describes the number of exposed complete MTSs. We find that the distributions of both the elongation rate and the MTS exposure time *t*_expo_ = (*L* – *l*_MTS_)/*k*_elong_ substantially differ between the two gene groups, with conditionally localized genes exhibiting more rapid elongation and shorter MTS exposure times (Fig. 2D; see Fig. S2 for similar distributions of conditional and constitutive gene elongation rates derived from (***Riba et al., 2019***)). These differences in MTS exposure kinetics between the two gene groups point towards a mechanism, thus far not part of our quantitative model, that would reduce the number of exposed MTSs (*β*), allowing for more variability in localization between the two groups. At the same time, this mechanism should have a greater effect in reducing MTS exposure time in conditionally localized genes, enabling reduced localization of this group at low mitochondrial volume fractions.

### Mitochondrial binding competence requires a maturation period

To reduce *β* and MTS exposure time, we introduce into our quantitative model a time delay between complete translation of the MTS and maturation of the MTS signal to become binding competent (Fig. 1C, *k*_MTS_ < ∞). This additional parameter is consistent with evidence that mitochondrially imported proteins require the recruitment of cytosolic chaperones to target them for recognition (***von Heijne, 1986***) and import by receptors on the mitochondrial surface (***Bykov et al., 2020***; ***Young et al., 2003***; ***Hoseini et al., 2016***). During MTS maturation, which could include autonomous folding or interaction with additional chaperone proteins (***Stein et al., 2019***), the MTS becomes capable of binding the mitochondrial surface.

In the model, MTS maturation is treated as a stochastic process with constant rate *k*_MTS_ corresponding to an average maturation time *τ*_MTS_ = 1/*k*_MTS_. This maturation period decreases the binding-competent exposure time uniformly across all mRNA, and decreases the numberof binding-competent MTS signals (i.e. lowers *β*) for all mRNA. The maturation period has the largest effect on short mRNAs with fast elongation, reducing their already short exposure times. Consequently, it is expected to have a larger effect on conditional versus constitutive genes.

The additional MTS maturation time does not alter the total time to translate an mRNA (*T*_total_ = *L*/*k*_elong_). The ribosome continues elongating during maturation, and is located at a downstream codon when the MTS becomes binding competent. The mean steady-state number of binding-competent MTSs per mRNA is

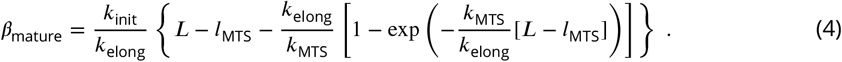

The mean time that each MTS is binding competent is

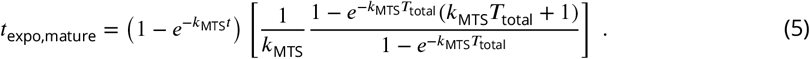

For mRNA localization to be sensitive to mitochondrial volume fraction, we expect the MTS exposure time to be shorter than the diffusive search times at low MVF (slow search, long search time) and longer than diffusive search times at high MVF (fast search, short search time). Such an intermediate exposure time will allow for high mitochondrial localization exclusively at high MVF.

The mean search time fora particle of diffusivity *D* to find a smaller absorbing cylinder of radius *r*_1_ when confined within a larger reflecting cylinder of radius *r*_2_ > *r*_1_ is (***Berg and Purcell, 1977***)

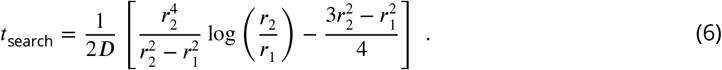

The smaller, absorbing radius *r*_1_ represents the cylinder sufficiently close to bind the mitochondrial surface, while *r*_2_ is the cylinder representing a typical distance that the diffusing particle must move through the cytoplasm to approach a different region of the mitochondrial network. As the mitochondrial volume fraction decreases, the radius *r*_2_ and the diffusive search time to find the mitochondrial surface *r*_search_ both increase.

To understand the impact of MTS maturation, we consider a typical conditional and constitutive mRNA from each group, using median translation rates and gene length. Figure 3A shows the exposure time *t*_expo,mature_ as the maturation time is varied. We find exposure times for a typical conditional gene to be intermediate between the high and low MVF diffusive search times when the maturation time is in the range *τ*_MTS_ = 10-100 seconds (Fig. 3A). By contrast, the typical constitutive gene maintains an exposure time that is higher than the diffusive search time for this parameter range.

**Figure 3.**
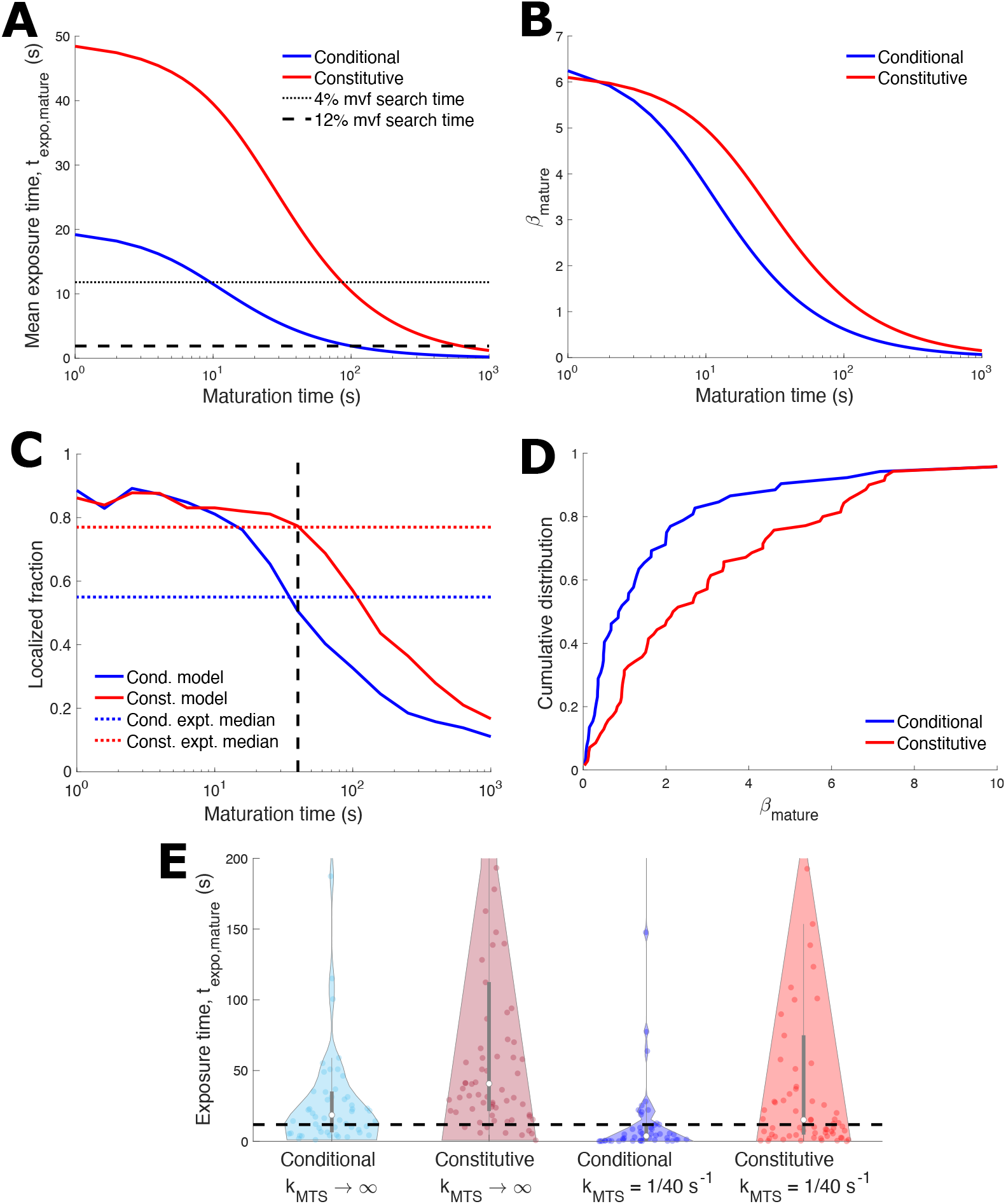
MTS binding-competence maturation time underlies distinct mitochondrial localization behavior of conditional and constitutive genes. (A) Mean exposure time of a binding-competent MTS before completing translation (Eq. 5) vs binding-competence maturation time. Data for median conditional (*L* = 393 aa, *k*_init_ = 0.3253 s^-1^, *k*_elong_ = 14.5086 s^-1^) and constitutive genes (*L* = 483 aa, *k*_init_ = 0.1259 s^-1^, *k*_elong_ = 7.7468 s^-1^) is shown. Horizontal dashed lines are the mean diffusive search times (Eq. 6) to reach binding range of mitochondria (region 1 in Fig. 1D). (B) *β*_mature_ (mean number of mature binding-competent MTS signals, Eq. 4) vs maturation time for median conditional and constitutive genes. (C) Mitochondrial localization (to region 1) vs maturation time for median conditional and constitutive genes with 4% MVF. Horizontal dotted lines indicate experimental localization medians. 40 second maturation time (vertical dashed line) allows model to match experimental localization for both conditional and constitutive genes. (D) Cumulative distribution of *β*_mature_ (mean mature MTS signals per mRNA) for conditional and constitutive genes. Steeper rise of conditional genes indicates more conditional genes have low *β* than constitutive genes; compare to Fig. 2A, which lacked MTS maturation time. (E) Violin plot showing model exposure times with 40-second MTS maturation and the instantaneous model without MTS maturation (*k*_MTS_ → ∞). 4% MVF. Median conditional exposure time with maturation is below the diffusive search time to find the binding region (horizontal dashed line, Eq. 6 for 4% MVF) while the other three medians are above this search time. For (C) – (E), the translation kinetics for each gene are estimated from experimental data (see Methods).

In addition to modulating the kinetics of binding competency, the maturation period decreases the expected number of functional MTS signals per mRNA, *β* (Fig. 3B). For the typical conditional gene, *β* decreases to approximately 1 for maturation times of 40 - 50 seconds, while *β* ≈ 2.5 for the typical constitutive gene in this range. The introduction of the MTS maturation time can thus selectively shift the expected number of functional MTS signals on conditional mRNA to the intermediate range (*β* ≈ 1) necessary to allow for MVF sensitivity in the localization behavior. Under the same conditions, the constitutive mRNA would maintain a high number of functional MTSs and thus should remain localized even at low MVF.

Figure 3C shows how the localization for the prototypical conditional and consitutive mRNA varies with the maturation time. For very rapid MTS maturation (*τ_MTS_* → 0), the MTS maturation model shows consistently high localization, as expected from the earlier model wherein the MTS became binding competent immediately upon translation. As the MTS maturation time increases and binding competency drops, both typical conditional and constitutive mRNA decrease their mitochondrial localization. However, the localization of the typical conditional mRNA begins to fall at approximately 10 seconds of maturation, while constitutive mRNA localization remains high until approximately 40 seconds of maturation. To provide a specific estimate of the maturation time, we determine the maturation times for which the model predicts the median experimental localization for conditional and constitutive genes (Fig. 3C, intersection of dotted lines and solid lines). A single value of *τ*_MTS_ ≈ 40 seconds yields a simultaneous accurate prediction for the localization of both groups (Fig. 3C, dashed).

Overall, the experimental data is consistent with a single gene-independent time-scale for MTS maturation. The stochastic model with a 40-second MTS maturation period was next applied to each of the conditional and constitutive mRNAs, for which translation parameters were calculated individually. With this maturation time, *β*_mature_ is substantially lower for conditional mRNA in comparison to constitutive mRNA (Fig. 3D).

For conditional mRNAs without the maturation period (*k*_MTS_ → ∞), the median MTS exposure time is greater than the diffusive search time (Fig. 3E, dashed black line). With a maturation time of *τ*_MTS_ = 40 s, the median conditional MTS exposure time decreases to be faster than diffusive search (Fig. 3E). In contrast, constitutive mRNAs retained a median MTS exposure time longer than the diffusive search time, both with and without the 40-second maturation period.

### Mitochondrial localization of conditional mRNAs is sensitive to inhibition of translational elongation and to mitochondrial volume fraction

Using the stochastic model with a 40-second MTS maturation period, we compute the localization of individual mRNAs in the constitutive and conditional groups, at a low mitochondrial volume fraction of 4%. Unlike the instantaneous model (with no MTS maturation delay), the localization of conditional genes is predicted to be significantly lower than that of constitutive genes (Fig. 4A). While introduction of this maturation time distinguishes the mitochondrial localization of conditional and constitutive gene groups (Fig. 4A vs Fig. 2B), other changes, such as to diffusivity, are unable to separate the two gene groups (Fig. S3).

**Figure 4.**
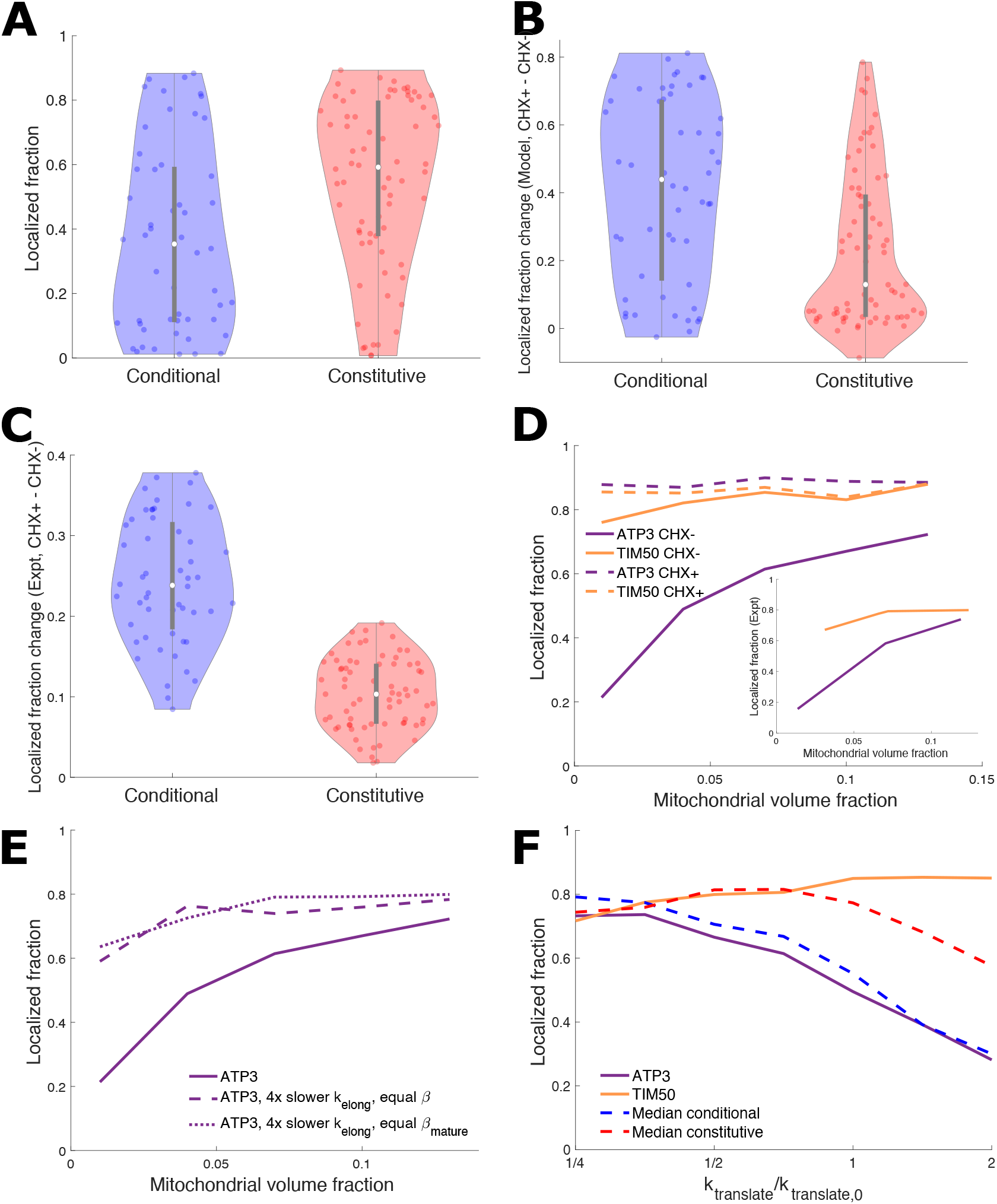
MTS maturation time distinguishes mRNA localization of conditional and constitutive genes. (A) Violin plots of mitochondrial localization of conditional and constitutive genes for model with 40-second maturation time; compare to Fig. 2B, which lacked MTS maturation time. p-value = 0.5% for two-sample Kolmogorov-Smirnov test for a difference between conditional and constitutive localization distributions. (B,C) Violin plots of localization increase upon cycloheximide application for model with 40-second MTS maturation time (B) and from experiment (C). (D) Mitochondrial localization for *ATP3* and *TIM50* vs MVF for model with 40-second MTS maturation time. Solid lines are CHX-, which closely corresponds to experimental data (***Tsuboi et al., 2020b***) (inset). Dashed lines are CHX+ model predictions, exhibiting large increase upon CHX application for *ATP3* and limited increase for *TIM50* (E) Comparing model mitochondrial localization results for *ATP3* to similar hypothetical construct gene with decreased elongation rate and initial rate selected to maintain either MTS number *β* or mature MTS number *β*_mature_. (F) Comparing model mitochondrial localization results for median conditional and constitutive genes, *ATP3*, and *TIM50* as both elongation and initiation rates (*k*_translate_) are varied. *k*_translate,0_ is the elongation or initiation rate for each of *ATP3*, *TIM50*, and median conditional and constitutive genes. For all panels, the translation kinetics for each gene are estimated from experimental data (see Methods). For (F), see Fig. 3 for median conditional and constitutive translation kinetics. (A), (B), and (F) use 4% MVF.

Furthermore, we use our model to predict localization in the presence of cycloheximide (CHX), which halts translation (***Schneider-Poetsch et al., 2010***). The localization difference in response to CHX application was used originally to define the constitutive and conditional groups (***Williams et al., 2014***). The effect of CHX is incorporated in the model by assuming that all mRNAs with an exposed MTS at the time of CHX application will be able to localize to the mitochondrial surface, since further translation will be halted by CHX. We therefore compute from our simulations the fraction of mRNAs that have at least one fully translated (but not necessarily mature) MTS, defining this as the localization fraction in the presence of CHX. The model predicts that conditional genes will have a substantial difference in localization upon application of CHX, while the difference for localization of constitutive genes will typically be much smaller (Fig. 4B). Qualitatively, this effect is similar to the observed difference in localization for experimental measurements with and without CHX (Fig. 4C).

The predicted mitochondrial localization of the two example mRNAs *ATP3* and *TIM50* is shown in Fig. 4D as a function of mitochondrial volume fraction. The model predicts *ATP3* localization is strongly sensitive to MVF, switching from below 30% at low MVF to above 70% localization at high MVF. By contrast, high localization of *TIM50* is predicted regardless of the MVF. The sensitivity of *ATP3* and insensitivity of *TIM50* localization to the MVF is consistent with experimental measurements indicating that *ATP3* exhibits switch-like localization under different metabolic conditions, while *TIM50* remains constitutively localized (***Tsuboi et al., 2020b***) (Fig. 4D, inset). Dashed lines in Fig. 4D show the predicted localization after CHX application, highlighting the difference in response to CHX between *ATP3* and *TIM50*.

The introduction of a delay period for MTS maturation both reduces the average number of binding-competent MTSs on each mRNA (lower *β*) and decreases the exposure time of each MTS. The latter effect results in faster switching between binding-competent and incompetent states for an mRNA. In the basic 4-state model, we saw that a steep sensitivity to the spatial region available for binding depends on having relatively rapid binding-state switching kinetics compared to the diffusion timescale (Fig. 1B). As shown in Fig. 3, the exposure time for conditional mRNAs is intermediate between the diffusive search times at high and low mitochondrial volume fractions. We therefore expect that the high rate of losing binding competence associated with the limited MTS exposure time to be critical for the switch-like response to mitochondrial volume fraction by *ATP3*.

As initiation rate can compensate for slowing translation elongation rates to maintain ribosome density (***Chu et al., 2014***; ***Kasari et al., 2019***), we consider hypothetical constructs which have the same average ribosome density (equal *β*) and mature MTS number (*β*_mature_) as *ATP3*, but 4-fold slower translational elongation rates. This results in slower switching kinetics, causing high localization and a loss of sensitivity to mitochondrial volume fraction (Fig. 4E). We also consider how translation rate adjustment could control mRNA localization while remaining at a fermentative mitochondrial volume fraction (4%). Localization substantially decreases with increasing elongation and initiation rates for the median conditional gene and *ATP3*, while localization is less responsive to increased translation rates for the median constitutive gene and *TIM50* (Fig. 4F). For responsive genes, translation rate modulation can adjust localization in a similar manner to mitochondrial volume fraction, with the potential for targeting of specific genes.

Overall, these results highlight the importance of translation kinetics, including both elongation rates and the maturation time of the MTS, in determining the ability of transcripts to localize to the mitochondrial surface. These kinetic parameters determine not only the equilibrated fraction of mRNAs that host a mature MTS but also the rate at which each mRNA switches between binding-competent and incompetent states. In order to achieve switch-like localization that varies with the mitochondrial volume fraction or CHX application, a transcript must exhibit an average of approximately one binding-competent MTS, with an exposure time that is intermediate between diffusive search times at low and high MVFs.

## Discussion

We have investigated, using quantitative physical modeling and analysis of yeast transcriptome data, the role of translation kinetics in controlling MTS-mediated localization of nuclear-encoded mRNA to mitochondria. Specifically, we explored how mRNA binding competence and association with the mitochondrial surface, across a range of cellular conditions, is governed by the interplay of timescales for translation and cytoplasmic diffusion. We compared two sets of mRNA: one that is localized conditionally, when mitochondrial volume is expanded or when translational elongation is halted by cycloheximide, and another that localizes constitutively regardless of these conditions. For these 52 conditional and 70 constutitive mRNA we estimated gene-specific translation kinetics to apply in the model. Our analysis indicates that these two sets of transcripts exhibit global differences in translation kinetics, and that these differences control mRNA localization to mitochondria by adjusting the number and duration of exposure for mitochondrial targeting sequences (MTSs) that are competent to bind to the mitochondrial surface.

It has previously been noticed when comparing mitochondrially localized versus non-localized yeast mRNAs, that localized mRNAs have features that reduce translation initiation and lower ribosome occupancy (***Poulsen et al., 2019***). This observation seemed counterintuitive as MTS exposure was thought to be important for the localization of many of these mRNAs and hence higher ribosome occupancy would be expected to enhance localization by increasing the number of exposed MTSs (***Sylvestre et al., 2003***; ***Williams et al., 2014***). While lower occupancy was proposed to drive mRNA localization through increased mRNA mobility of a poorly loaded mRNA (***Poulsen et al., 2019***), our results propose an alternative explanation. We show that translational parameters which yield a moderate number of approximately 1 – 2 binding competent ribosomes (via associated MTSs) per mRNA nevertheless allow robust localization under physiological conditions. Furthermore, this model occupancy allows for localization levels to be steeply sensitive to mitochondrial volume fraction, enabling transcript localization to be modulated by the MVF during changes to nutrient conditions and the metabolic mode. By constrast, transcripts with a high occupancy are expected to remain constitutively localized to mitochondria, regardless of the metabolic state of the cell. Thus, tuning of translational kinetics allows for differential response of transcript localization under varying nutrient conditions without the need for additional signaling pathways.

Translation kinetics can vary widely vary between genes, with greater than 100-fold variation in mRNA translation initiation rates and approximately 20-fold variation of elongation rates in yeast (***Riba et al., 2019***). Translation duration can be further impacted by the length of the coding sequence. Constitutively localized mRNAs are on average longer and have slower translation elongation than conditionally localized mRNAs. Cis regulators of translation elongation rates include mRNA features such as codon usage, codon context, and secondary structures (***Gebauer and Hentze, 2004***; ***Espah Borujeni and Salis, 2016***). For the constitutively localized mRNA *TIM50* it was previously found that a stretch of proline residues, which are known to slow ribosome elongation, were necessary to maximize mRNA localization of this mRNA to the mitochondria (***Tsuboi et al., 2020b***).

To investigate the role of these varied parameters, we first explore an abstracted four-state model, wherein each transcript can be near or far from the mitochondrial surface and competent or not for binding to the mitochondria. This model shows that increasing the equilibrium fraction of time in the binding-competent state is indeed expected to enhance mitochondrial localization. Furthermore, the simplified model demonstrates that in order for transcript localization to be sensitive to the fraction of space where binding is possible (i.e., the mitochondrial volume fraction), the kinetics of switching in and out of the binding-compentent state must be relatively rapid compared to the kinetics for spatial movement.

We then proceed to develop a more detailed model that explicitly incorporates translational initiation and elongation, the formation of an MTS that enables mitochondrial binding, and diffusive search for the mitochondrial surface. This model confirms that tuning of the translation parameters can substantially alter mitochondrial localization, but only in a regime where the ribosome occupancy of the transcripts is relatively low. Surprisingly, plugging physiological parameters into this instantaneous model resulted in the prediction that all mRNA transcripts studied would be highly localized to mitocondria in all conditions. In other words, the physiological parameters appeared to be in a regime where most transcripts had multiple binding-competent MTS sequences with long exposure time, resulting in global localization.

Motivated by differences in transcript length and elongation rate between constitutive and conditional gene groups, we incorporated an MTS maturation period into the model, driving the system into a paremeter regime with lower numbers of binding-competent MTSs and shorter MTS exposure times, particularly for the more rapidly elongating and shorter conditional transcripts. Although we are unable to directly attribute this maturation period to a particular process, it aligns with other observations related to mitochondrial protein import. It is known that mitochondria targeting sequences mediate interactions with mitochondrial recognition machinery, namely TOM22 and TOM20 subunits of the translocase of the outer membrane (TOM) complex, and are necessary for efficient protein import into the mitochondria (***Backes et al., 2018***). The folding process for some proteins that must be recognized and imported into mitochondria occurs on a timescale that competes with translocation (***Mukhopadhyay et al., 2004***; ***Regev-Rudzki et al., 2008***). Furthermore, the formation of a secondary structure has been shown to be required for import of MTS-bearing proteins into mitochondria (***Waltner et al., 1996***). Together, these observations suggest the MTS is likely to require time to mature prior to becoming fully competent. Slowed translation has been suggested as providing an opportunity for proteins to fold, implying the MTS maturation time may also be regulated by translation kintetics (***Zhao et al., 2021b***).

In addition, molecular chaperones such as Hsp70 and Hsp90 are important for the delivery and recognition of the mitochondrial preproteins to the Tom70 receptor (***Deshaies et al., 1988***; ***Young et al., 2003***). Hsp70 expression levels have been found to have a direct effect on mRNA localization to the mitochondria (***Eliyahu et al., 2012***). STI1 is another cochaperone of Hsp70 and Hsp90 chaperones that plays a role in recognizing mitochondrial preproteins and mediates targeting to the mitochondria (***Hoseini et al., 2016***). All of these data point to the need for a delay time between MTS translation and its maturation into a binding-competent state, via either autonomous folding or association with a chaperone, before it can be optimally recognized by the surface of the mitochondria.

Upon incorporation of a uniform (gene-independent) 40-second MTS maturation time into the model, we found that many genes fell into a parameter regime with only a few mature, binding-competent MTS sequences per transcript, and with intermediate exposure times for those sequences. This single choice of the maturation time made it possible to simultaneously match the expected localization of prototypical constructs representing both the constitutive and conditional gene groups. This choice of parameter yielded a mature MTS exposure time in the conditional gene that was longer than the diffusive search time at high mitochondrial volume fraction, yet shorter than the search time at low volume fraction. Consequently, the model with an MTS maturation time could adequately predict the decreased localization of conditional genes under metabolic conditions with low MVF, while genes in the constitutive group were localized regardless of the MVF.

Notably, conditional localization in our model required not only a modest number of mature MTS per transcript (*β*_mature_ ≈ 1) but also relatively fast translational initiation and elongation kinetics (short exposure times compared to diffusive search). This result demonstrates the out-of-equilibrium nature of the localization process, wherein localization is dictated by the kinetic rates themselves rather than their ratios or the equilibrated fraction of transcripts in different states. This feature arises due to broken detailed balance (***Gnesotto et al., 2018***) in the kinetic scheme illustrated in Fig. 1A, wherein binding-competent transcripts bind irreversibly to the mitochondrial surface and can be dislodged only by the completion of the energy-consuming translation process. Subcellular localization of mRNA can thus be added to the extensive list of biomolecular processes wherein the tools of non-equilibrium statistical mechanics elucidate the relevant physical parameters governing system behavior (***Murugan et al., 2012***; ***Gladrow et al., 2016***; ***Maza et al., 2019***; ***Brown and Sivak, 2019***; ***Fang and Wang, 2020***; ***Mogre et al., 2020***).

We approximate mitochondrial geometry in the cell as a central mitochondria cylinder concentrically within a larger cell cylinder. This approximation represents a scenario where an mRNA that diffuses too far from one mitochondrial tubule has become closer to another tubule. This approximation neglects mitochondrial fusion and fission dynamics, as well as the non-uniform distribution of mitochondria to the yeast cell periphery (***Viana et al., 2020***). A non-uniform mitochondrial distribution could introduce longer diffusion times for some mRNA trajectories before a mitochondrial tubule encounter, likely further decreasing the already weak localization of some mRNA at low MVF. Further exploration of the role of spatial mitochondrial distribution in mRNA localization is left as a fruitful area for further study.

From the perspective of biological function, it remains unclear why some mitochondrial mRNAs localize conditionally under different metabolic conditions, while others remain constitutively localized. Both types contain an MTS (***Williams et al., 2014***; ***Elstner et al., 2009***) and code for proteins rich in hydrophobic residues that are susceptible to misfolding and aggregation in the cytosolic space (***von Heijne, 1986***). One reason for the differential localization may center on the altered function of mitochondria from fermentative to respiratory conditions. ATP synthase, the linchpin of the mitochondrial OXPHOS metabolic process, is comprised of subunits of both prokaryotic and eukaryotic origin (***Rühle and Leister, 2015***). Interestingly, all but one of the prokaryotic-origin subunits are conditionally localized to the mitochondria (***Tsuboi et al., 2020b***). As mitochondrial mRNA localization has been found to be sufficient to upregulate protein synthesis (***Gehrke et al., 2015***; ***Tsuboi et al., 2020b***) we posit that conditional or switch-like localization behavior is a post-transcriptional regulation mechanism of protein synthesis that is sensitive to mitochondrial growth and metabolic state. In particular, this mechanism can act globally, altering expression levels for a large set of transcripts, even without the involvement for specific signaling pathways to adjust protein synthesis in response to metabolic state.

Furthermore, we propose that the effects of a respiratory metabolic state, which increases mitochondrial volume fraction and decreases the mRNA diffusion search time, can be mimicked through global translation elongation inhibition by pushing MTS signal dynamics into a much slower regime than mRNA diffusive search, potentially altering mitochondrial composition. This hints at translation elongation inhibition as an avenue or tool for toggling metabolic modes within the cell. Similar means of post-transcriptional regulation may take place in mammalian cells as genome-wide mRNA localization measurements to the mitochondria have found a class of mRNAs that are constitutively localized while others are found to become localized after CHX administration (***Fazal et al., 2019***).

Our results link the nonequlibrium physics governing localization of transiently binding-competent mRNA and the observed differential response of transcript groups that localize to mitochondria under varying metabolic conditions. The general principles established here, including the importance of translation kinetics and transport timescales to the organelle surface, apply broadly to cellular systems that rely on a peptide targeting sequence for co-translational localization of proteins. For example the localization of mRNAs encoding secretory proteins to the surface of the endoplasmic reticulum (ER) through interactions between the signal recognition sequence on the nascent peptide, the signal recognition particle that binds it, and receptors on the ER surface, may well be governed by analogous principles (***Zhang and Shan, 2012***; ***Zhao et al., 2021a***). By coupling together quantitative physical models and analysis of measured translational parameters for the yeast transcriptome, this work provides general insight on the mechanisms by which a cell regulates co-translational localization of proteins to their target organelles.

## Methods

### Simplified discrete-state model

Figure 1A describes a minimal model for mRNA localization with four discrete states: sticky and close (*S*_N_), sticky and far (*S*_F_), not sticky and close (*U*_N_), and not sticky and far (*U*_F_). mRNA can transition between these states with rates *k*_R_, *k*_L_, *k*_U_, and *k*_S_, as shown in Fig. 1A. These transitions are mathematically described by

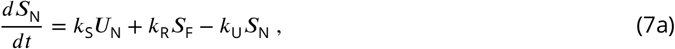

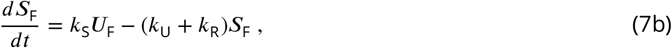

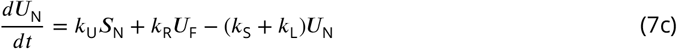

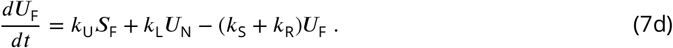

Note that there is no direct transition from *S*_N_ to *S*_F_ because if an mRNA is bound to the mitochondria it cannot leave the mitochondrial vicinity. Setting all derivatives in Eqs. 7 to zero, the steady-state solution is

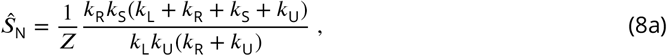

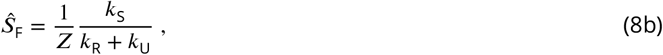

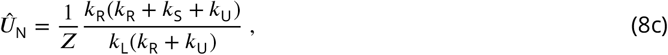

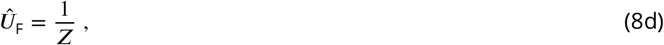

with

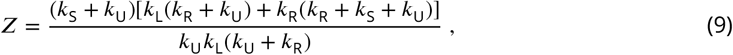

for state probabilities 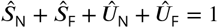.

In the regime where mRNA transport is much faster than the binding-competence switching rate (*k*_R_, *k*_L_ ≫ *k*_U_, *k*_S_), the near fraction is

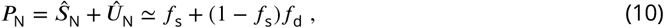

where *f*_s_ = *k*_S_/(*k*_S_ + *k*_U_) and *f*_d_ = *k*_R_/(*k*_R_ + *k*_L_). In the opposite regime, where mRNA transport is much slower than the binding-competence switching rate (*k*_R_, *k*_L_ ≪ *k*_U_, *k*_S_), the near fraction is

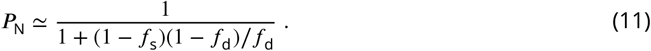

### Stochastic simulation with translation and diffusion

We use stochastic simulations to determine mitochondrial mRNA localization and fraction of time spent in the binding-competent state. Individual (non-interacting) mRNA molecules are simulated from synthesis in the nucleus to decay in the cytosol.

#### mRNA synthesis, translation, and MTS binding competence

The mRNA simulation begins after exit from the nucleus, as experiments can fluorescently label and track mRNA once synthesized in the nucleus. The time spent by mRNA in the nucleus is a normally-distributed time period with mean 60 sand standard deviation of 30 seconds (if a negative time is selected, the distribution is resampled until a positive time is yielded). After nuclear exit, the mRNA begins simulated translation and diffusion through the cytosol.

Each mRNA has *L* codons. Ribosomes arrive and initiate translation with rate *k*_init_ if the first codon is not occupied. Each ribosome on an mRNA moves forward to the next codon at rate *k*_elong_ if the next codon is not occupied. A ribosome on the *L*’th (final) codon completes translation at rate *k*_elong_, leaving the final codon unoccupied. mRNA decay at a rate *k*_decay_ once in the cytosol. The parameters *k*_init_, *k*_elong_, and *L* are varied to represent different genes (see below for the calculation of *k*_init_ and *k*_elong_ for particular genes). The mRNA decay rate is set to *k*_decay_ = (1/600) s^-1^. Stochastic translation trajectories are generated using the Gillespie algorithm (***Gillespie, 1977, 2007***).

We applied two models of mRNA gaining mitochondrial binding competence through mitochon-drial targeting sequence (MTS) translation. For the instantaneous model, mRNA are competent to bind mitochondria if there is a least one ribosome at or past codon *l*_MTS_ = 100. For the maturation model, once a ribosome reaches *l*_MTS_ = 100, the ribosome will gain competence to bind the mRNA to a mitochodrion at a rate *k*_MTS_. This rate *k*_MTS_ is included in the Gillespie algorithm, to select when a ribosome will confer binding competence.

#### Diffusion

The cell volume is defined as concentric cylinders (Fig. 1D). The central cylinder is the mitochondria, which is maintained at a radius *r*_m_ = 350 nm. The radius *R* of the outer cylinder is selected to establish a desired mitochondrial volume fraction. A typical yeast cell volume is *V* = 42 *μ*m^3^. We assume that 80% of this volume is not occupied by the nucleus and vacuole, and thus available to mitochondria, the cytosol, and other cell components. Thus, the mitochondrial volume fraction in the simulation 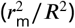 is set equal to 0.8*f_m_* where *f_m_* is the reported volume fraction. Specifically, we set 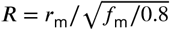. We note that this outer radius represents not the size of the cell as a whole, but rather the typical separation between non-proximal tubes within the mitochondrial network. A particle that hits the boundary of this outer cylinder would then begin to approach either the same or another mitochondrial network tube. We thus treat the outer cylinder as a reflecting boundary.

The simulation uses a propagator approach to sample the transitions of the mRNA between concentric regions around the mitochondrion, analogous to previous approaches used to simulate the dynamics of DNA-binding proteins (***Koslover et al., 2011***) and diffusing organelles (***Mogre and Koslover, 2018***). The closest region (region 1), for radial distances *r*_m_ < *r* < *r_a_* = *r*_m_ + 25 nm, is sufficiently close for a binding-competent mRNA to bind a mitochondrion. mRNA within the intermediate cylindrical shell (region 2), with *r_a_* < *r* < *r_b_* = *r*_m_ + 250 nm, are sufficiently close to the mitochondrion that they appear close in diffraction-limited imaging but are not sufficiently close to be able to bind. The last cylindrical shell (region 3), for *r_b_* < *r* < *R*, represents the cell region where an mRNA would not be near any mitochondria.

In the simulations, region 1 is treated as a cylinder with an absorbing boundary at *r_a_* + *є*. A particle that first enters the region is placed at initial position *r_a_* – *є* and the first passage time to the absorbing boundary is sampled from the appropriate Green’s function for radially symmetric diffusion in a cylindrical domain (***Özisik, 1993***). Region 2 is treated as a hollow cylinder with absorbing boundaries at *r_a_* – *є* and *r_b_* + *є*. Particles that enter region 2 from region 1 start at position *r_a_* + *є* and those that enter from region 3 start at *r_b_* – *є*. Region 3 is a hollow cylinder with absorbing boundary at *r_b_* – *є* and reflecting boundary at *R*. Particles that enter region 3 from region 2 start at position *r_b_* + *є*. The buffer width to prevent very short time-steps at the region boundaries is set to *є* = 10 nm. If the sampled transition time for leaving a region occurs before the next translation process selected by the Gillespie algorithm, the mRNA changes regions and the translation state transition times are then resampled. mRNAs that first exit the nucleus are placed at position *r* = *R*.

Binding-competent mRNA in region 1 are unable to leave this region, because they are bound to the mitochondrion. When a binding-competent mRNA in this region loses binding competence, the mRNA is given a random radial position within *r*_m_ < *r* < *r_a_*, with the probability of the radial position proportional to *r*.

#### Localization measures

We use two types of localization measures, corresponding to different experimental measurements. One measure considers an mRNA localized to mitochondria if the mRNA is close enough to bind (*r*_m_ < *r* < *r*_m_ + 25 nm). This measure corresponds to experiments that chemically bind nearby mRNA to mitochondria to determine the fraction localized. The other measure considers an mRNA localized if the mRNA is close enough that with diffraction-limited imaging the mRNA appears next to the mitochondria (*r*_m_ < *r* < *r*_m_ + 250 nm). While quantitatively distinct, these measures do not lead to qualitatively different results.

#### Ensemble averaging

For each localization measurement shown in our results, we simulate 50 mRNA trajectories from synthesis to decay, with each trajectory having a lifetime (including time spent in the nucleus) and a fraction of that lifetime spent mitochondrially localized. The ensemble average is calculated by weighting the fraction localized of each trajectory by the trajectory lifetime,

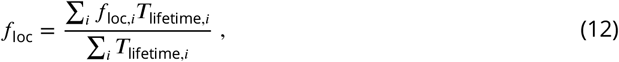

where *f*_loc,*i*_ is the fraction of trajectory *i* spent mitochondrially localized and *T*_lifetime,*i*_ is the mRNA lifetime for trajectory *i*. The probability that an mRNA will be included in a localization measurement, through either experimental localization measurement technique, is proportional to the lifetime of the mRNA.

### Calculation of translation rates

We assume that each mRNA produces proteins at a rate *k*_init_, so that the cell produces a particular protein at a rate *N*_mRNA_*k*_init_, where *N*_mRNA_ is the number of mRNA for a gene. For a steady state number of proteins, protein production must be balanced by protein decay. We assume that the primary mode of effective protein decay is cell division, such that each protein has an effective lifetime equal to a typical yeast division time of *T*_lifetime_ = 90 minutes. The steady-state translation initiation rate is then taken as

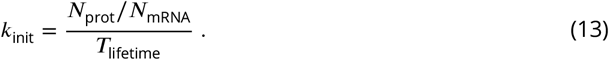

Protein per mRNA data (***Couvillion et al., 2016***; ***Morgenstern et al., 2017***) provides relative, rather than absolute, numbers for the number of proteins in a cell per mRNA of the same gene. Accordingly, we can rewrite our expression for *k*_init_ as,

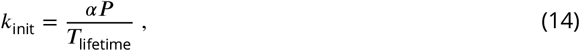

where *P* is the protein per mRNA measurement (***Couvillion et al., 2016***; ***Morgenstern et al., 2017***), and *α* is the proportionality constant. To calibrate, we use the gene *TIM50* as a standard, as there are available measurements of *N*_prol_ = 4095 (***Morgenstern et al., 2017***) and *N*_mRNA,TIM50_ = 6 (***Tsuboi et al., 2020b***). From Eq. 13, *k*_init,TIM50_ = 0.1264 s^-1^, and with *P*_TIM50_ = 15.12 and from Eq. 14 gives *α* = 45.14. With *α* and *P*, we estimate *k*_init_ across genes.

The steady-state number of ribosomes *N*_ribo_ on an mRNA balances ribosome addition to the mRNA at rate *k*_init_ and removal at rate *k*_elong_*N*_ribo_/*L*, such that *k*_elong_ = *k*_init_*L*/*N*_ribo_. Ribosome occupancy *R* (***Zid and O’Shea, 2014***) is proportional to the ribosome density *N*_ribo_/*L*. We can thus write,

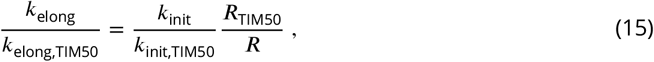

and apply *k*_elong,TIM50_ = 4 aa/s (***Riba et al., 2019***) to estimate *k*_elong_ across genes.

### Calculating MTS exposure time and mature MTS numbers per mRNA

In this section Eqs. 4 and 5 are derived.

We assume MTS maturation is a Poisson process, i.e. with constant rate *k*_MTS_. The probability that an MTS has not yet matured at time *t* after its translation is *I*(*t*) = *e*^*^−k^MTS^t^*^. After the MTS has been translated, the ribosome completes translation after a mean time *t_max_* = (*L* – *l*_MTS_)/*k*_elong_. For an MTS that matures before the ribosome terminates translation, the mean waiting time *t*_wait_ from MTS translation to maturity is

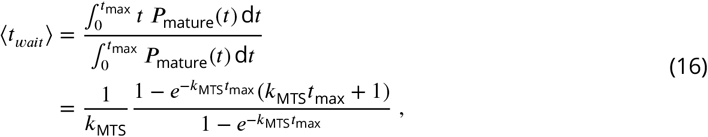

where *P*_mature_ = *k*_MTS_*I*(*t*).

A fraction *I*(*t*_max_) of translated MTS regions do not mature before translation termination, so the mean time that a mature MTS is exposed on the mRNA is

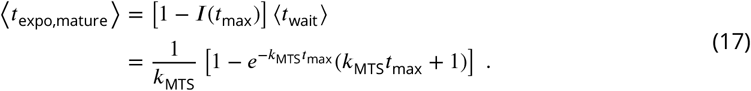

The number of mature MTSs per mRNA, *β*_mature_, is related to the mean number of ribosomes per mRNA codon, *ρ*_ribo_ = *k*_init_/*k*_elong_. The probability that an MTS is mature at time *t* after ribosome initiation is 1 – *I*(*t*). The ribosome reaches codon *x* beyond its initiation point at time *t*(*x*) = *x*/*k*_elong_ Integrating over the codons beyond the MTS region,

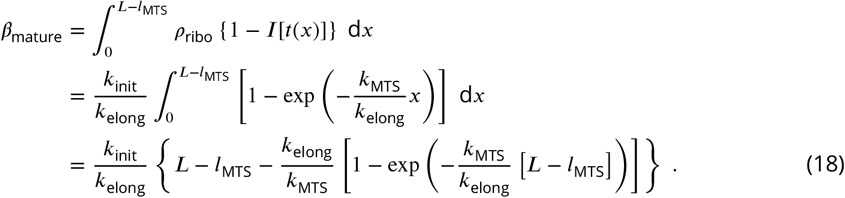

## Acknowledgements

We thank TTsuboi, M Viana, and RSubramaniam for helpful discussions and feedback on the paper. AIB was supported by a Natural Sciences and Engineering Research Council of Canada Discovery Grant 2021-03431 and start-up funds provided by the Ryerson University Faculty of Science. EFK was supported by NSF grant #2034482, and a Cottrell Scholar Award from the Research Corporation for Science Advancement. BMZ was supported by NIH grant R35GM128798.

## Conflict of Interests

The authors declare that they have no conflict of interest.

## Notes

### Competing Interest Statement

The authors have declared no competing interest.

### Summary of Updates

Added to Acknowledgements and added a Conflict of Interest statement.

