## Supplemental Figures for "Mitochondrial mRNA localization is governed by translation kinetics and spatial transport"

### Supplemental Information for "Mitochondrial mRNA localization is governed by translation kinetics and spatial transport"

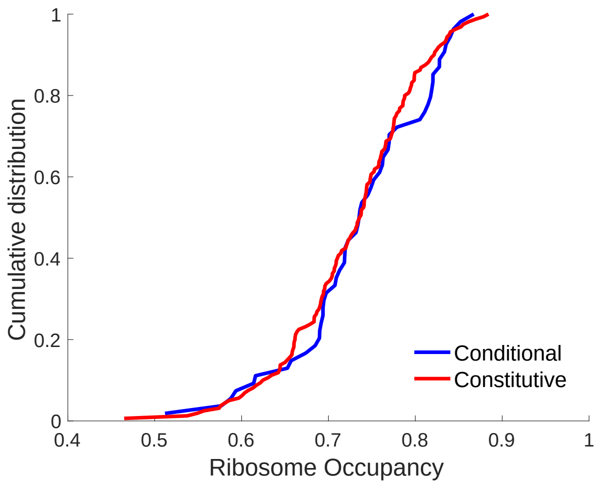


FIG. S1. Cumulative distribution of conditional and constitutive mRNA genes vs ribosome occupancy (lines indicate fraction of genes with given ribosome occupancy or less). Ribosome occupancy from Arava et al [1]. $n_{\text{conditional}} = 54$ and $n_{\text{constitutive}} = 160$.

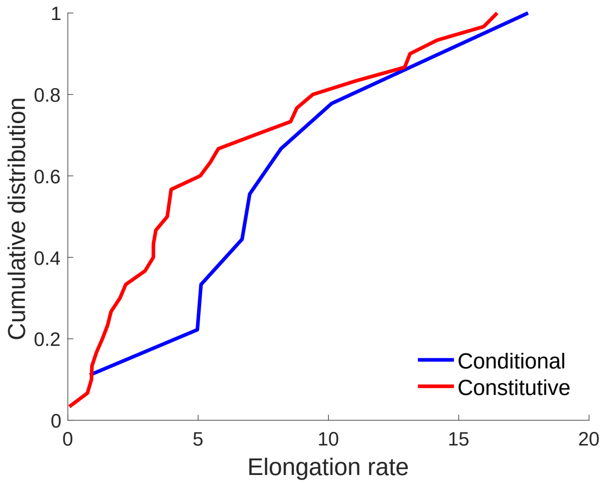


FIG. S2. Cumulative distribution of conditional and constitutive genes vs elongation rates (lines indicate fraction of genes with given elongation rate or less). Elongation rates calculated with data from and as described in Riba et al [2], with elongation rate equal to protein synthesis rate divided by ribosome density. $n_{\text{conditional}} = 9$ and $n_{\text{constitutive}} = 30$.

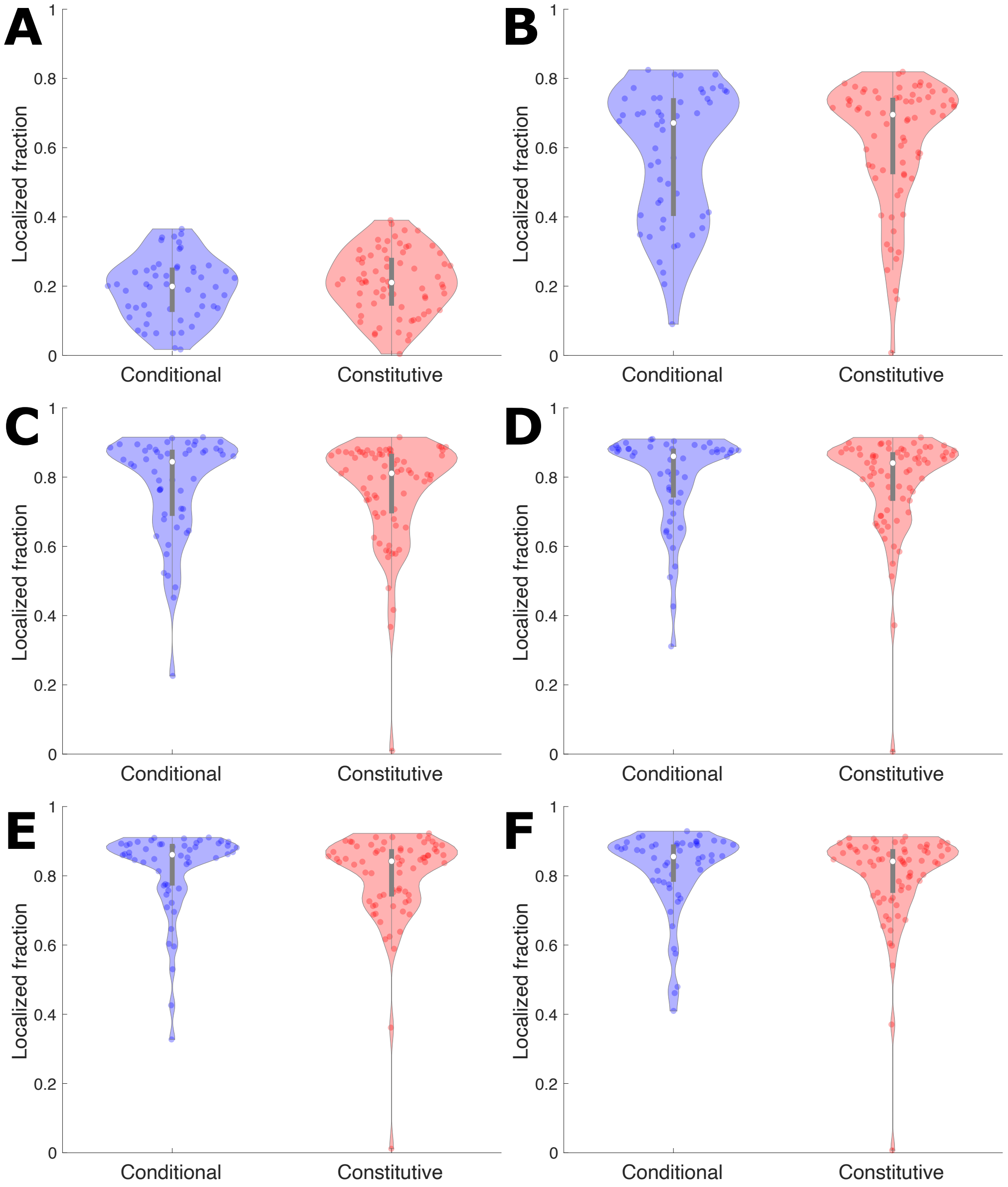


FIG. S3. Violin plot showing mRNA localization fraction of individual genes with instantaneous model (no maturation delay) with translation kinetics for each gene estimated from experimental data (see Methods) and 4% MVF. (A) is with mRNA diffusivity $D = 0.001\ \mu\text{m}^2/\text{s}$, (B) with $D = 0.01\ \mu\text{m}^2/\text{s}$, (C) with $D = 0.1\ \mu\text{m}^2/\text{s}$, (D) with $D = 0.2\ \mu\text{m}^2/\text{s}$, (E) with $D = 0.5\ \mu\text{m}^2/\text{s}$, and (F) with $D = 1\ \mu\text{m}^2/\text{s}$.

[1] Yoav Arava, Yulei Wang, John D Storey, Chih Long Liu, Patrick O Brown, and Daniel Herschlag. Genome-wide analysis of mrna translation profiles in saccharomyces cerevisiae. *Proceedings of the National Academy of Sciences*, 100(7):3889–3894, 2003.

[2] Andrea Riba, Noemi Di Nanni, Nitish Mittal, Erik Arhné, Alexander Schmidt, and Mihaela Zavolan. Protein synthesis rates and ribosome occupancies reveal determinants of translation elongation rates. *Proceedings of the National Academy of Sciences*, 116(30):15023–15032, 2019.
